# Bone marrow transplantation rescues monocyte recruitment defect and improves cystic fibrosis in mice

**DOI:** 10.1101/570135

**Authors:** Zhichao Fan, Jacqueline Miller, Rana Herro, Erik Ehinger, Douglas J. Conrad, Zbigniew Mikulski, Yanfang Peipei Zhu, Paola M. Marcovecchio, Catherine C. Hedrick, Klaus Ley

**Affiliations:** Division of Inflammation Biology, La Jolla Institute for Immunology, 9420 Athena Circle Drive, La Jolla, California 92037, USA; Division of Immune Regulation, La Jolla Institute for Immunology, 9420 Athena Circle Drive, La Jolla, California 92037, USA; Microscopy and Histology Core Facility, La Jolla Institute for Immunology, 9420 Athena Circle Drive, La Jolla, California 92037, USA; Division of Pulmonary, Critical Care and Sleep Medicine, Department of Medicine, University of California San Diego, 9500 Gilman Drive, La Jolla, California 92093, USA; Department of Bioengineering, University of California San Diego, 9500 Gilman Drive, La Jolla, California 92093, USA

## Abstract

In this study, we demonstrate that correcting the monocyte adhesion defect in CFTR^ΔF508^ mice (CF mice) by bone marrow transplantation significantly improved survival and reduced inflammation.

**Abstract:** Cystic fibrosis (CF) is an inherited life-threatening disease accompanied by repeated lung infections and multi-organ inflammation that affects tens of thousands of people worldwide. The causative gene, cystic fibrosis transmembrane conductance regulator (CFTR), is mutated in CF patients. Monocytes from CF patients show a deficiency in integrin activation and adhesion. Since monocytes play critical roles in controlling infections, defective monocyte function may contribute to CF progression. In this study, we demonstrate that monocytes from CFTR^ΔF508^ mice (CF mice) show defective adhesion under flow. Transplanting CF mice with wild-type bone marrow after sublethal irradiation replaced most (60-80%) CF monocytes with wild-type monocytes, significantly improved survival, and reduced inflammation. Wild-type/CF mixed bone marrow chimeras directly demonstrated defective CF monocyte recruitment to the bronchoalveolar lavage and the intestinal lamina propria in vivo. Our findings show that providing wild-type monocytes by bone marrow transfer rescues gastrointestinal (GI) mortality in CF mice, suggesting that wild-type bone marrow stem cells might mitigate CF inflammation.

## Introduction

Cystic fibrosis (CF) is one of the most common monogenic diseases (more than 70,000 cases worldwide, ~1,000 new cases each year) *(1)*, which is caused by variations in the cystic fibrosis transmembrane conductance regulator (CFTR) gene. CFTR is an anion-conducting transmembrane channel that is very important in mucus function and the ion homeostasis of cells(2). More than 2000 gene variants have been identified, which are grouped into six types of mutations*(2)*. In humans, CF disease is dominated by lung and pancreas pathologies. The dominant pathology in the lung is inflammation caused by failure to clear microorganisms and the generation of a toxic pro-inflammatory microenvironment *(2–4).* CFTR dysfunction in the pancreas results in pancreatic insufficiency in most CF patients *(2)*.

CFTR mutations result in mucus dysfunction *(5)* and impaired mucocilliary clearance in sino-pulmonary tissue. The resulting viscous mucus becomes infected with pathogens, such as *Pseudomonas aeruginosa *(6–9)*, Staphylococcus aureus *(10, 11)*, Haemophilus influenza (12)*, and *Aspergillus* species *(13).* These recurrent infections greatly exacerbate the symptoms of CF *(4, 5, 14)* and impact patient quality of life. Current clinical management of CF includes prophylactic (flucloxacillin) and therapeutic antibiotics (nebulized tobramycin, colistin, and aztreonam), DNAse (Dornase alfa), CFTR potentiators and correctors (Ivacaftor, Lumacaftor), and mucus thinners (Hypertonic saline) *(15)*. Although progress has been made *(16)*, there is currently no cure for CF.

There are several mouse models of CF *(17, 18)*, in which the homologues of common human CFTR mutations have been knocked into the mouse CFTR locus. Here, we use the CFTR^ΔF508^ mouse *(19)*, a widely used and well-accepted mouse model of CF *(20, 21).* The pathology in this model is dominated by intestinal pathology, which exhibits mucus accumulation in the crypts of Lieberkühn, goblet cell hypertrophy, hyperplasia, and eosinophilic concretion in the crypts, resulting in constipation and rectal prolapse *(22).* Death ensues when constipation leads to an ileus.

A recent study reported that mutations of CFTR commonly found in CF patients cause a profound monocyte adhesion defect *(23, 24).* The authors demonstrated expression of CFTR in human monocytes and found a significant defect of their integrin activation and static adhesion.

A similar defect of static adhesion was reported in CFTR^ΔF508^ mouse monocytes, resulting in a ~60% (human) or ~40% (mouse) loss of monocyte static adhesion. Activation of the three main monocyte integrins *(25)*, β_2_ (α_L_β_2_, α_M_β_2_) and α_4_ (α_4_β_1_), was found to be defective. Remarkably, this defect is restricted to monocytes, with no defect in neutrophils and a minor defect (~20-30%) in lymphocytes. These features and the molecular mechanism make it different from the known leukocyte adhesion deficiencies LAD I, II and III *(26)*, suggesting that CF is associated with a new leukocyte adhesion deficiency, LAD IV *(24)*.

Monocytes are important guardians of epithelial surfaces. They serve by directly secreting cytokines *(27)* and killing pathogens *(28, 29)* and can differentiate into macrophages and inflammatory dendritic cells *(30)*. Macrophages clear apoptotic debris (efferocytosis), secrete cytokines for tissue homeostasis and survey the environment *(31)*. When macrophages detect pathogens or danger signals, they directly phagocytose and kill bacteria *(29, 32).* They also produce many inflammatory cytokines and chemokines that orchestrate the recruitment of other immune and inflammatory cells *(27)*, which are involved in both the inflammatory antimicrobial response, wound healing and fibrosis *(33)*. Dendritic cells migrate to the draining lymph nodes, where they present antigens to CD4 and CD8 T cells and thus control the adaptive immune response.

Given the expression of CFTR in monocytes *(23)*, and given the monocyte adhesion defect, we reasoned that monocyte recruitment to mucosal sites might be defective in CFTR^ΔF508^ mice (CF mice), resulting in weakened host defense and increased pathogen burden. Here, we report that this is indeed the case. We reasoned that correcting CFTR in monocytes might improve CF disease burden and clinical outcomes. To test this, we performed bone marrow transplantation (BMT) to generate mice that were defective in CFTR in hematopoietic cells, non-hematopoietic cells, both, or neither. We observed improvement of CF symptoms and longer survival in mice that received wild-type (WT) bone marrow (BM). Mechanistically, we show that CF monocytes have a severe recruitment defect to the intestinal lamina propria and the bronchoalveolar space. Since it is now possible to correct genetic defects in HSCs *(34, 35)*, autologous transfer of engineered HSCs or allogeneic BMT would seem a plausible potential treatment for patients with CF.

## Results

### Leukocytosis in CF mice and adhesion defect of CF monocytes

First, we confirmed that CF mice died spontaneously under SPF conditions. The first mice succumbed at 23 days of age, with half of all deaths occurring by 40 days and ~91% CF mice succumbing by 59 days (Fig. 1A). Like wild-type (CFTR^WT/WT^) mice, heterozygous mice (CFTR^ΔF508/WT^) survived for at least 110 days. At the same age (paired comparison of littermates), CF mice had significantly lower body weights (Fig. 1B) and significantly elevated blood leukocyte counts (Fig. 1C). Monocytes, lymphocytes, and neutrophils were all elevated (Fig. 1D-F).

**Fig. 1.**
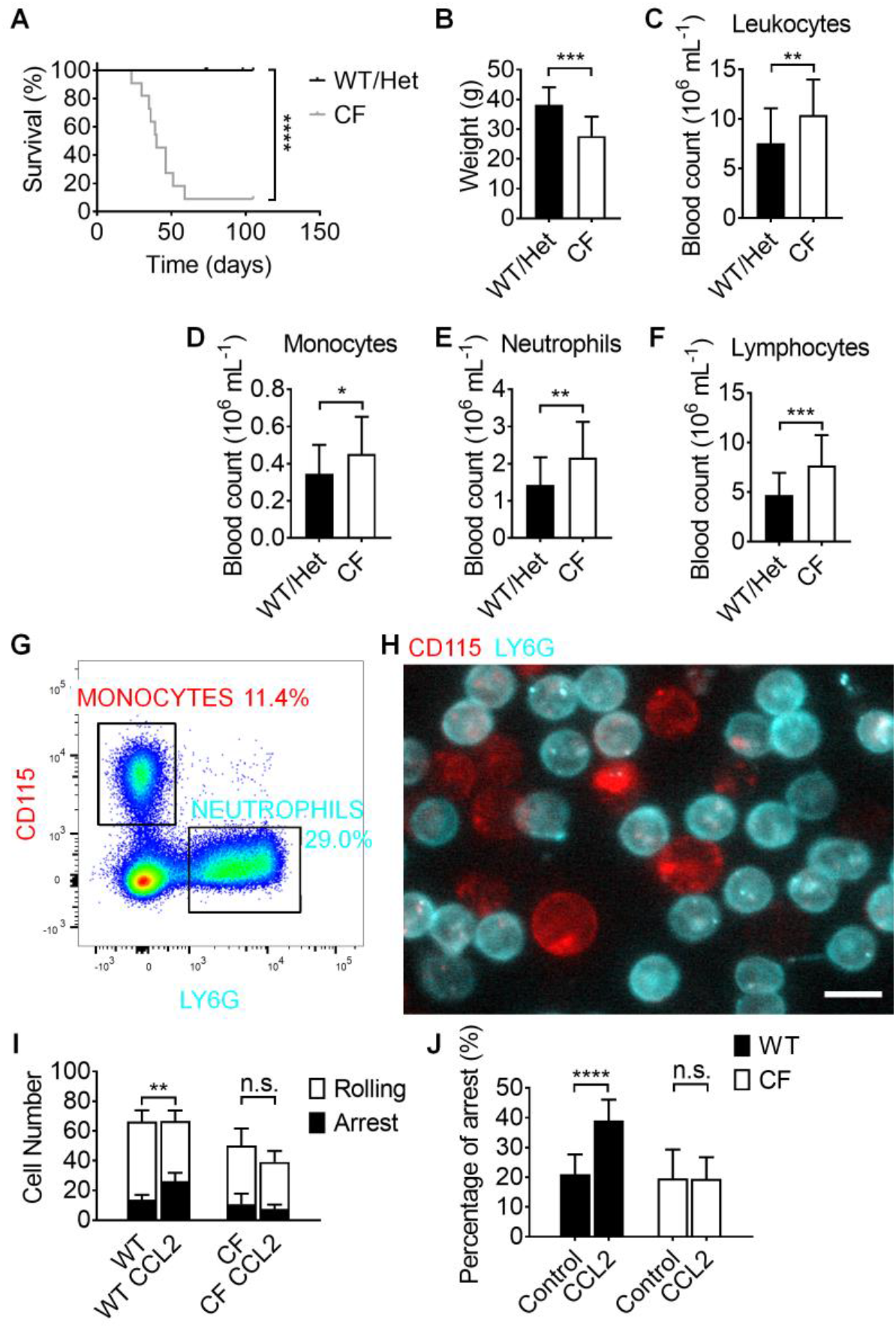
Leukocytosis and monocyte adhesion deficiency in CFTR^ΔF508^ mice. **(A-B)** the survival curve (A, n=21 WT and 11 CFTR^ΔF508^) and weight (B, n=12 each,) of CFTR^ΔF508^ and WT mice. **(C-F)** Hematology for total leukocytes (C, n=47 WT and 26 CFTR^ΔF508^), monocytes (D, n=47 WT and 26 CFTR^ΔF508^), neutrophils (E, n=35 WT and 17 CFTR^ΔF508^), and lymphocytes (F, n=35 WT and 17 CFTR^ΔF508^) in CFTR^ΔF508^ mice compared to WT controls. **(G-H)** Monocytes and neutrophils in bone marrow cells are distinguished by staining of CD115 and Ly6G staining, respectively, in flow cytometry (G) and by fluorescence microscopy in a microfluidic flow chamber (H). Both monocytes (red) and neutrophils (cyan) adhered on the substrate of P-selectin/ICAM-1 at a shear stress of 6 dyn·cm^−2^ (H). The scale bar is 10 μm. **(I-J)** Number of rolling (white) and arrested monocytes at rest and after infusion of CCL2 (100 ng·mL^−1^) (I). The percentage of arrested as a fraction of all monocytes in a field of view (J) for WT and CF monocytes with or without the stimulation of CCL2. n.s. p>0.05, *p≤0.05, **p≤0.01, ***p≤0.001, ****p≤0.0001 by Gehan-Breslow-Wilcoxon test (A) and student T-test (B-F,I-J).

Monocytes express CD115 (Colony-stimulating factor 1 receptor, CSF1R) while Ly6G identifies neutrophils in mouse BM (Fig. 1G). Using monoclonal antibodies to CD115 (red) and Ly6G (cyan) allowed us to visualize monocytes and neutrophils in flow chambers coated with P-selectin (to support rolling*(36, 37)*) and ICAM-1 (to support arrest*(37)*) (Fig. 1H). In these experiments, arrest was triggered by infusing CCL2 (Chemokine C-C motif ligand 2, also knowns as monocyte chemoattractant protein-1, MCP-1) at a concentration of 100 ng·mL^−1^. The same number of WT or CF BM cells (5×10^7^ mL^−1^) were perfused in the flow chambers. Rolling and arrest were monitored by epifluorescence microscopy. The total number of cells (monocytes plus neutrophils) per field-of-view (FOV) was the same in flow chambers perfused with WT or CF BM cells. The total number of monocytes per FOV was slightly less in the flow chambers perfused with CF than WT BM cells (Fig. 1I). The number of arrested WT, but not CF monocytes, doubled in response to CCL2 (Fig. 1I). Thus, we conclude that CCL2-induced arrest under flow is completely abolished in CF monocytes.

### Transplanting WT BM reliefs the disease in CF mice

Histological assessment of the small intestine revealed the previously described increased crypt depth and villus height in CF mice (Fig. 2A). For both parameters, the differences between CF and WT mice were significant (Fig. 2B, C). CF mice also showed more mucus accumulation in the crypts of Lieberkühn than WT mice (Fig. 2D), resulting in a significantly larger percentage of crypts containing mucus (Fig. 2E).

**Fig. 2.**
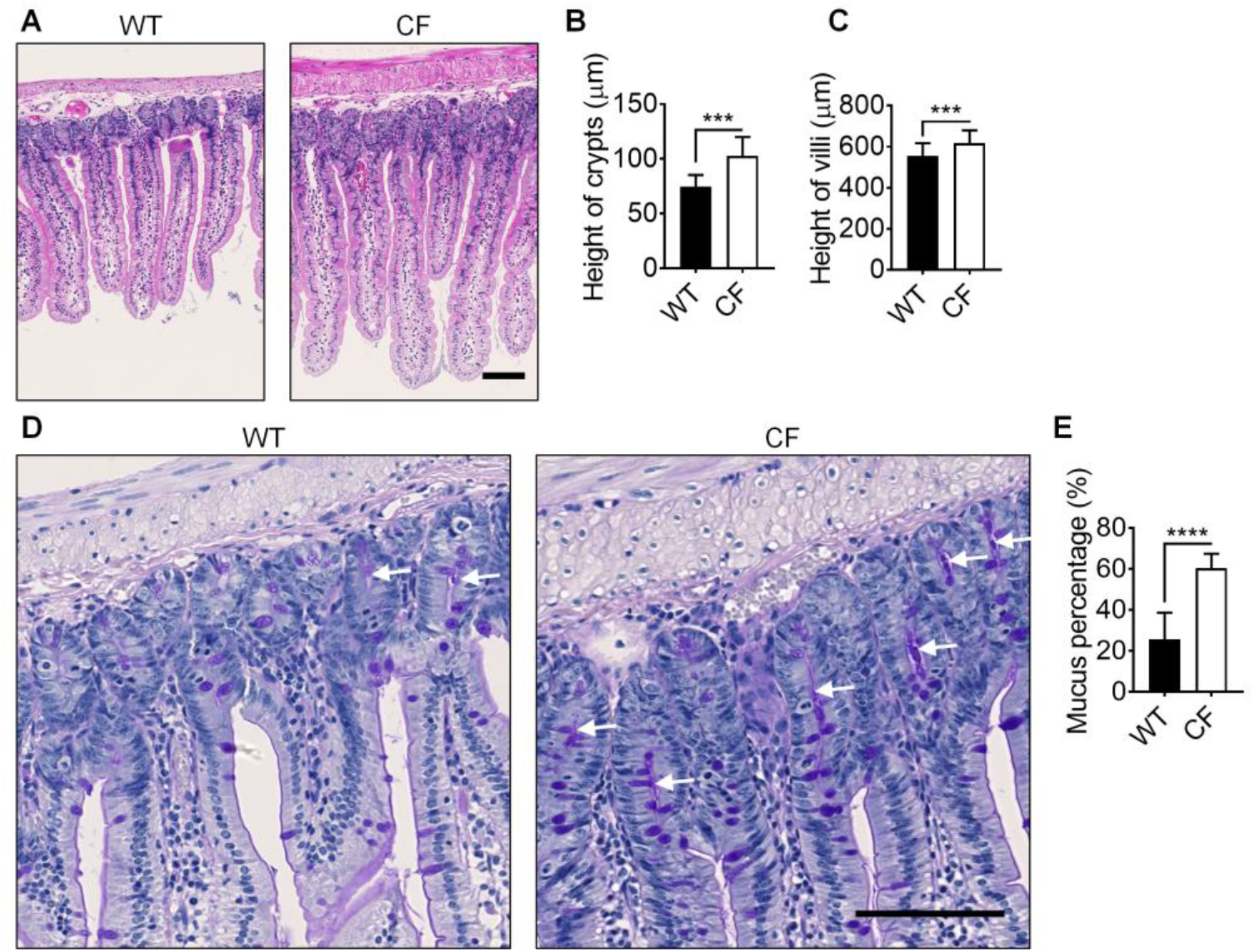
Intestinal inflammation in CFTR^ΔF508^ mice. **(A)** Typical H&E staining images of intestines from both WT and CFTR^ΔF508^ mice. **(B-C)** Crypt (B) and villi (C) height of small intestine from CFTR^ΔF508^ mice compared to WT controls. n=9 each. **(D)** Typical PAS staining images of intestines from both WT and CFTR^ΔF508^ mice. White arrows indicate mucus in crypts. **(E)** Percentage of intestinal crypts that contain mucus in them. n=27. ***p≤0.001, ****p≤0.0001 in the student T-test. Scale bars are 100 μm.

Next, we asked whether reconstituting CFTR in monocytes was sufficient to improve the health of CF mice. To this end, we conducted BMT by injecting sterile WT or CF (control) BM cells into sub-lethally (700 rads) irradiated CF recipient mice. Reconstituting CF mice with WT BM resulted in 60-80% WT monocytes (Table S1) and significantly prolonged their survival (Fig. 3A). The median survival time increased from 34 to 99 days, and 33% of the mice survived for up to 225 days. Interestingly, all mice that survived to 110 days (80 days after BMT) remained alive for the entire experiment. Pathologic evaluation of the bone marrow-transplanted mice showed that reconstituting CF mice with WT BM restored the crypt height defect (Fig. 3B, C), but not the villus height defect (Fig. 3D). This treatment also reduced the mucus percentage as assessed by histology (Fig. E, F).

**Fig. 3.**
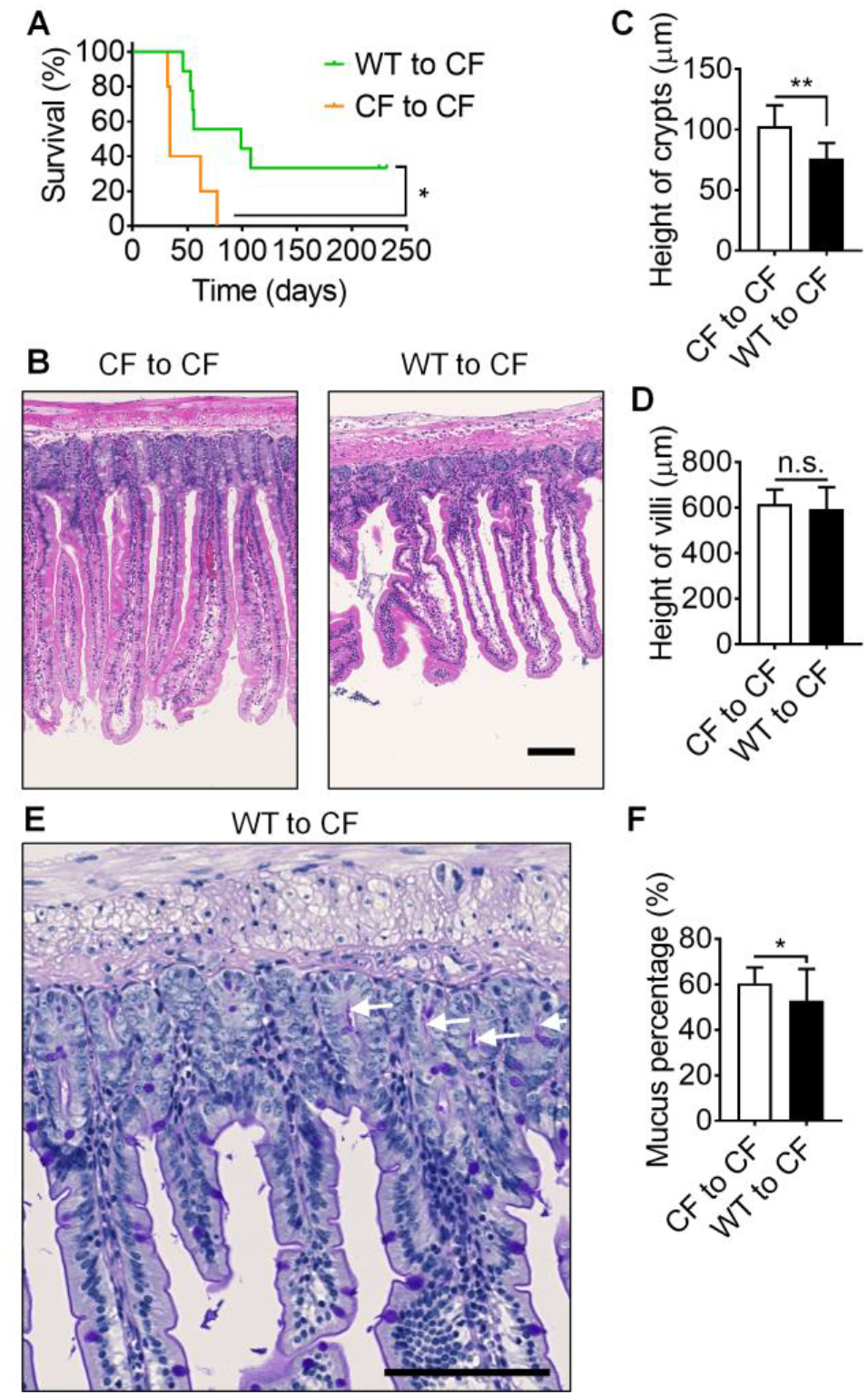
WT Bone marrow transplantation rescues the survival and intestine inflammation in CFTR^ΔF508^ mice. **(A)** the survival curve of CFTR^ΔF508^ mice (700 rads) transplanted with WT (green curve, n=9;) or CFTR^ΔF508^ (orange curve, n=5) BM. **(B)** Typical H&E staining images of intestines from CFTR^ΔF508^ mice transplanted with CFTR^ΔF508^ or WT BM. **(C-D)** Quantification of crypts (C) and villi (D) height of intestines from CFTR^ΔF508^ mice transplanted with CFTR^ΔF508^ or WT BM. n=9. **(E)** A typical PAS staining image of intestine from a CFTR^ΔF508^ mouse transplanted with WT BM. White arrows indicate mucus in crypts. **(F)** Percentage of intestine crypts that contain mucus in it. CFTR^ΔF508^ mice are transplanted with WT or CFTR^ΔF508^ BM. n=27. n.s. p>0.05, *p≤0.05, **p≤0.01in the Gehan-Breslow-Wilcoxon test (A) and student t-test (C-D,F). Scale bars are 100 μm.

### Transplanting CF BM induces the disease in WT mice

In a reverse approach, we tested whether the CFTR-dependent monocyte adhesion defect was sufficient to confer disease to WT mice. We reconstituted lethally (1100 rads) irradiated WT mice with CF or WT (control) BM (Fig. 4A). In these mice, almost all hematopoietic cells are donor-derived (Table S1), but the epithelial cells are host-derived and thus retain intact, functional CFTR. All WT to WT control mice survived for the entire duration of the experiment (232 days), but half of the mice that were reconstituted with CF BM succumbed by 41 days.

**Fig. 4.**
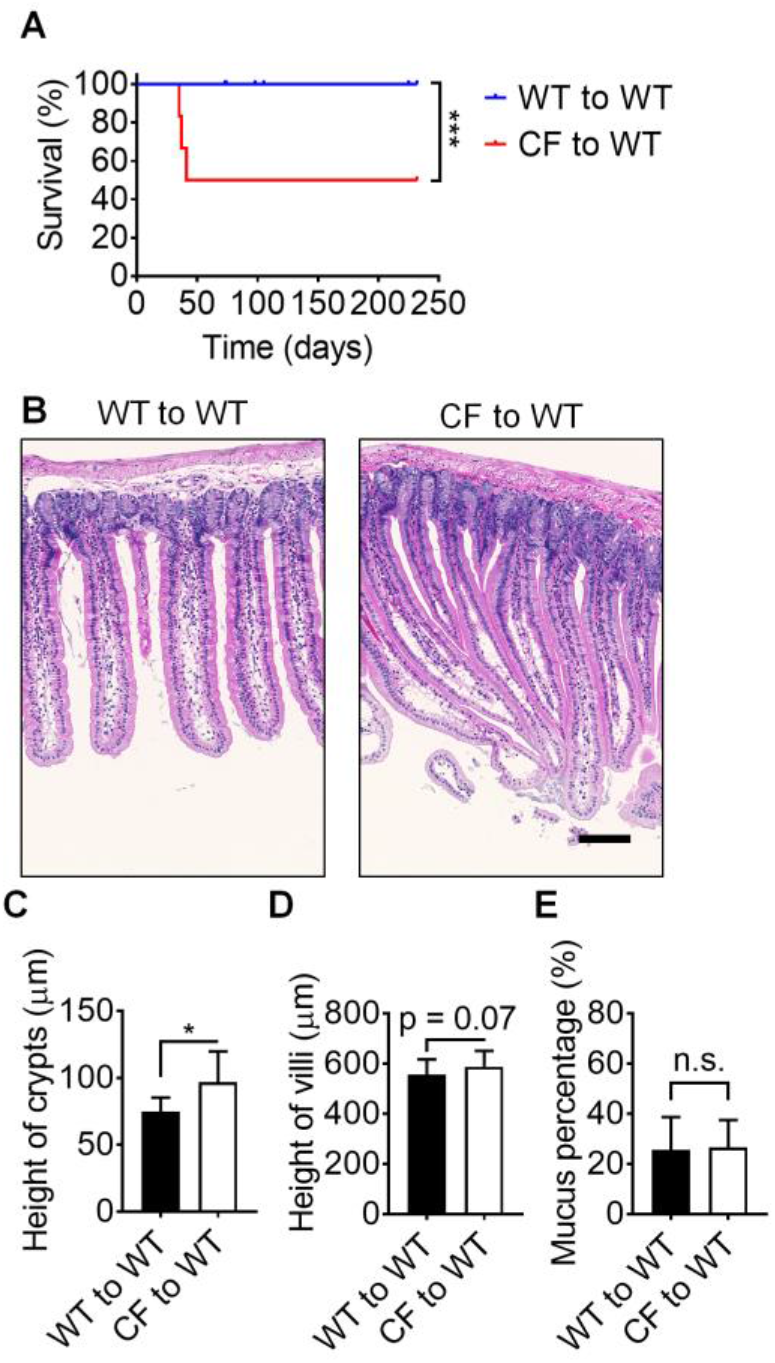
CFTR^ΔF508^ Bone marrow transplantation induces the disease in WT mice. **(A)** the survival curve of WT mice (1100 rads) transplanted with WT (blue curve, n=21) or CFTR^ΔF508^ (red curve, n=6) BM. **(B)** Typical H&E staining images of intestines from WT mice transplanted with CFTR^ΔF508^ or WT BM. **(C-D)** Quantification of crypts (C) and villi (D) height of intestines from WT mice transplanted with CFTR^ΔF508^ or WT BM. n=9. **(E)** Percentage of intestine crypts that contain mucus in it. WT mice are transplanted with WT or CFTR^ΔF508^ BM. n=27. n.s. p>0.05, *p≤0.05, ***p≤0.001 in the Gehan-Breslow-Wilcoxon test (A) and student t-test (C-E). Scale bars are 100 μm.

Reconstituting WT mice with CF BM induced the crypt height defect (Fig. 4B, C), but not villus height or mucus percentage (Fig. D, E). Thus, we conclude that CFTR^ΔF508^ in hematopoietic cells (by BMT of CFTR^ΔF508^ cells) significantly induces CF disease in WT mice. The CFTR-dependent monocyte adhesion defect is both necessary and sufficient to drive CF disease in mice. The reconstitution efficiency of BM in mice upon the 1100 rads lethally irradiation was >90% (Table S1).

### Recruitment defect of CF monocytes in vivo

Having shown the monocyte arrest defect under flow, we next asked whether this arrest defect translated into an in vivo monocyte recruitment defect of CF monocytes. To stringently investigate the recruitment of monocytes to the intestine and lung, we used mixed BM chimeras*(37)*. In these experiments, lethally irradiated WT mice were reconstituted with both WT and CF BM (Fig. 5A) at an approximately 1:1 ratio, tracked by congenic markers (CD45.1/CD45.2). Both monocytes and macrophages were identified by flow cytometry (gating scheme in fig. S1–3). In unchallenged mice, there was no difference in the number of WT and CF monocytes in the lung (Fig. 5B), but the lungs contained significantly more WT than CF macrophages (Fig. 5C), suggesting that the monocyte-to-macrophage differentiation may be defective in CF monocytes. To directly test monocyte recruitment in vivo, we challenged mixed CF/WT bone marrow chimeric mice with intranasal CCL2. Significantly more WT than CF monocytes appeared in the bronchoalveolar lavage fluid (BAL), demonstrating defective CF monocyte recruitment in vivo (Fig. 5D). This was also true for the Ly6C-hi subset of monocytes (Fig. 5E). In the small intestinal lamina propria, the findings were similar: we found no difference between WT and CF monocytes in unchallenged mice (Fig. 5F), but more WT than CF macrophages in lamina propria leukocytes (LPL, Fig. 5G). Taken together, these findings indicate that the CF monocyte defect manifests in both elicited monocyte recruitment and in monocyte to macrophage differentiation.

**Fig. 5.**
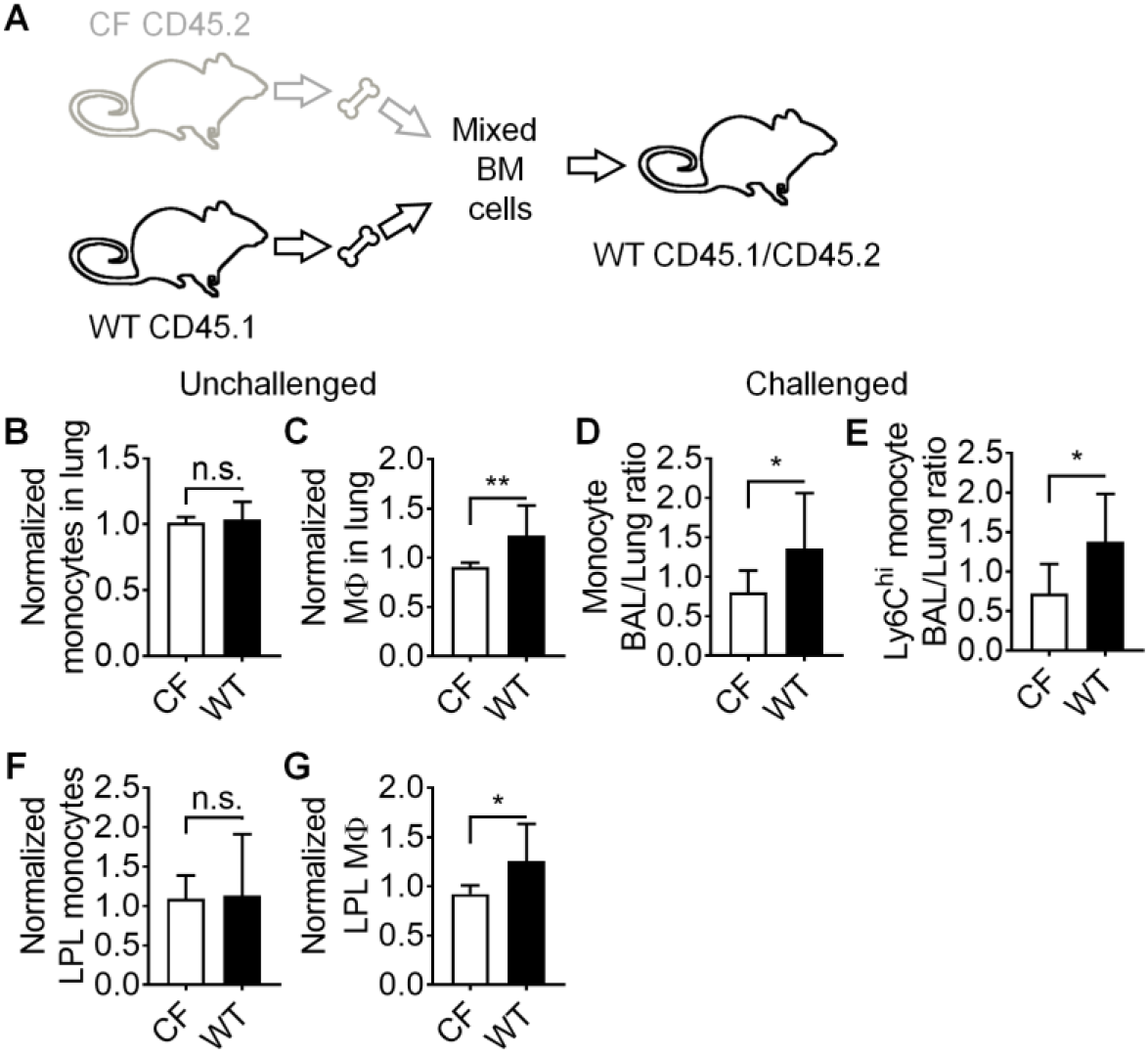
Monocyte recruitment in mixed bone marrow chimeras. **(A)** Schematic showing the establishment of CFTR^ΔF508^/WT mixed bone marrow chimeras. **(B-C)** CFTR^ΔF508^ and WT monocytes (CD115^+^CD11b^+^CD11c^−^F4/80^−^, B) and macrophages (F4/80+CD11b^+^, C) in the lungs of unchallenged mixed chimeric mice as assessed by flow cytometry, see fig. S1–3 for gating scheme. **(D-E)** Mixed chimeric mice were challenged with CCL2 (intranasal 100 ng/mice, 12 hours before the harvest) and the BAL/lung monocyte ratio (D) or Ly6C^hi^ BAL/lung monocyte ratio (E) was calculated for CFTR^ΔF508^ and WT monocytes. **(F-G)** The small intestinal lamina propria leukocytes (LPL) of mixed chimeric mice were harvested assessed for monocytes (CD11b^+^CD11c^−^F4/80^−^Ly6G^−^TCR^−^Cd19^−^NK1.1^−^, F) and macrophages (F4/80+, G) by flow cytometry. n=10 for B-C, F-G and14 for D-E. n.s. p>0.05, *p≤0.05, **p≤0.01 in the student T-test.

## DISCUSSION

This study reports mouse experiments demonstrating that BMT rescues the CF phenotype in the CFTR^ΔF508^ mouse model, suggesting that BMT could mitigate some aspects of CF disease by restoring monocyte/macrophage function and mucosal host defense. The survival and intestinal inflammation of CF mice were both improved by the hematological reconstitution using WT BM. We show that CF monocytes fail to adhere under flow, show reduced recruitment in response to CCL2, and contribute less to macrophages in the lung and intestinal lamina propria than WT monocytes. We show that the CFTR^ΔF508^ in monocytes is both necessary and sufficient for full manifestation of CF disease.

Besides the recruitment defect, CF monocytes may have other functional defects when challenged by inflammation and infections. Upon LPS stimulation, monocytes from CF patients have been reported to show reduced inflammatory cytokine expression (lower TNFα, lower IL-6, lower IL-12, lower IL-23) and increased production of anti-inflammatory IL-10 compared to WT controls *(38)*. Taken together with our findings, this suggests complex defects encompassing arrest, recruitment, cytokine production and differentiation into macrophages. The common denominator of these defects may be defective β_2_ integrin activation in CF monocytes, because β2 integrins are known to be involved in many myeloid cell functions *(39)*.

Whether or not CF monocytes show defective phagocytosis remains controversial. Complement-mediated phagocytosis of C3b-coated beads and *Pseudomonas aeruginosa* has been reported to be deficient in blood monocytes from CF patients *(40)*. In another study, the phagocytosis of *Escherichia coli* was increased in CF monocytes *(38)*. These findings suggest that some but not all phagocytosis mechanisms are affected by dysfunctional CFTR. Antigen presentation to CD4 T cells by MHC-II is decreased in CF monocytes *(38)*. A recent study showed that wild-type CFTR associates with PTEN, triggering the anti-inflammatory PI3K/Akt pathway in response to TLR4. This helps enhance *Pseudomonas aeruginosa* clearance *(7)*, suggesting a mechanism that may fail in CF patients.

CFTR is expressed in the monocyte, but not the neutrophil plasma membrane *(40)*. Consequently, CFTR deficiency did not affect the adhesion of human neutrophils *(23)*, which are the most abundant leukocytes in human blood and play essential roles in immune defense *(37)*. However, CFTR mRNA is expressed in human neutrophils *(47)*. CFTR protein is found expressed in the phagolysosome membrane of neutrophils, which affects bacterial killing, degranulation and disease outcome *(41–46).* Thus, correcting CFTR by BMT may also restore the function of neutrophils and improve CF by promoting bacterial killing.

Four previous studies used BMT to ask whether stem cells existed that may restore CFTR function in lung *(47–49)* or gut *(49, 50)* epithelial cells. These studies showed that hematopoietic stem cells and multipotent mesenchymal stromal cells have the ability to home to the lung or intestine and differentiate into epithelial cells. A mild but significant rescue (~10%) of epithelial electrophysiology was shown in distal colons of CF mice, which have a decrease of Forskolin-induced transepithelial current compared to WT controls *(49, 50).* However, the epithelial reconstitution was incomplete, even under the optimized conditions *(47)*. Optimized WT to CF BMT improved bacterial clearance and survival after infection with *Pseudomonas aeruginosa (47).* In light of the present findings, it seems more likely that the benefit observed in *(47)* was due to the functional rescue of myeloid cell function rather than epithelial cell function.

Myeloid cell recruitment to the lung is a multi-step process, where the myeloid cells first leave the blood compartment to appear in the lung interstitium and then must negotiate the epithelial layer to appear in the bronchoalveolar lavage *(51)*. Our present findings suggest that the monocyte recruitment defect is localized to the transepithelial and not the transendothelial migration. The recruitment defect is not detected under homeostatic conditions but appears when mice are challenged with CCL2. CF patients likely are in a permanently challenged condition because of the bacterial load in the bronchoalveolar mucus. The mechanisms leading to a significantly reduced percentage of CF monocyte-derived macrophages in the lung and the intestinal lamina propria require further study.

Taken together, we show that restoration of WT monocytes is sufficient to improve health and survival in CF mice. Since many CF patients are diagnosed at a young age, and since BMT is more successful in children than in adults, this is an appealing strategy *(52)*. An antibody-based safer approach for BMT has been developed *(53)*, which might be suitable for testing our proposed CF treatment. Unlike in malignant diseases, lethal irradiation is not required and partial restoration with 60-80% WT monocytes is sufficient. Moreover, it is now possible to correct genetic defects in HSCs *(34, 35)* by gene therapy approaches, so that autologous transfer of engineered HSCs also may become a practical treatment for patients with CF.

## MATERIALS AND METHODS

### Reagents

Recombinant mouse P-selectin-Fc, ICAM-1-Fc and CCL2 were purchased from R&D Systems. Casein blocking buffer and Penicillin-Streptomycin solution were purchased from Thermo Fisher Scientific. Roswell Park Memorial Institute (RPMI) medium 1640 without phenol red, phosphate-buffered saline (PBS) without Ca^2+^ and Mg^2+^, Hanks’ balanced salt solution (HBSS) with phenol red without Ca^2+^ and Mg^2+^ were purchased from Gibco. Fetal bovine serum (FBS) and human serum albumin (HSA) were purchased from Gemini Bio Products. Type VIII collagenase, Dnase, and N-Acetyl-L-cysteine I were purchased from Sigma-Aldrich. Phycoerythrin (PE) or PE-Cyanine 7 (Cy7)-conjugated anti-CD115 antibody (clone AFS98), Alexa Fluor 647 (AF647) or AF700-conjugated anti-Ly6G antibody (clone 1A8), Brilliant Violet 785 (BV785)-conjugated anti-CD11b antibody (clone M1/70), BV605-conjugated anti-CD11c antibody (clone N418), BV570-conjugated anti-Ly6C antibody (clone HK1.4), BV711-conjugated anti-TCRβ antibody (clone H57-597), Allophycocyanin (APC)-conjugated anti-CD117 antibody (clone 2B8), APC-Cy7-conjugated anti-CD19 antibody (clone 6D5), PE-conjugated anti-Ly6A/E antibody (clone D7), BV650-conjugated anti-NK1.1 antibody (clone PK136), and PE-conjugated anti-F4/80 antibody (clone BM8) were purchased from Biolegend. Ghost Dye™ Violet 510 was purchased from Tonbo biosciences. Fluorescein (FITC)-conjugated anti-CD45.1 antibody (clone A20) was purchased from BD Biosciences. Peridinin Chlorophyll Protein Complex (PerCP)-Cy5.5-conjugated anti-CD45.2 antibody (clone 104), PE-eFluor610-conjugated anti-CD4 antibody (clone GK1.5), eFluor450-conjugated anti-CD8a antibody (clone 53-6.7) were purchased from eBioscience. Red blood cell (RBC) lysis buffer was purchased from Invitrogen.

### Mice

C57BL/6J wild-type mice (000664; JAX), DsRed mice (006051; JAX; wild-type C57BL/6J background), CD45.1 mice (002014; JAX; wild-type C57BL/6J background), and CFTR^Δ508^ mice (002515; JAX; C57BL/6J background) were obtained from the Jackson Laboratory. CD45.1/CD45.2 mice were breed by C57BL/6J wild-type and CD45.1 mice. Mice were fed a standard rodent chow diet and were housed in microisolator cages in a pathogen-free facility. Mice were euthanized by CO_2_ inhalation. All experiments followed guidelines of the La Jolla Institute for Allergy and Immunology Animal Care and Use Committee, and approval for use of rodents was obtained from the La Jolla Institute for Allergy and Immunology according to criteria outlined in the Guide for the Care and Use of Laboratory Animals from the National Institutes of Health.

### BMT

WT/Het or CFTR^Δ508^ Recipients (C57BL/6J background littermates, male or female, one month old) were irradiated in two doses of 550 or 350 rads each (for a total of 1,100 or 700rads) four hours apart (RS-2000 X-Ray irradiator, Rad Source). Bone marrow cells from both femurs and tibias of donor mice (WT/Het or CFTR^Δ508^ mice, sex matched, one month old) were collected under sterile conditions. Bones were centrifuged for the collection of marrow, unfractionated bone marrow cells were washed, resuspended in PBS, injected retro-orbitally into the lethally irradiated mice (one donor to five recipients). Recipient mice were housed in a barrier facility under pathogen-free conditions before and after bone marrow transplantation. After bone marrow transplantation, mice were provided autoclaved acidified water with antibiotics (trimethoprim-sulfamethoxazole) and were fed autoclaved food. The survival of mice was monitored. Mice were used for further experiments eight weeks after bone marrow reconstitution. In some experiment, DsRed WT mice were used as recipients or donors to test the reconstructive rate of BMT by flow cytometry.

In the mixed chimeric BMT, CD45.1/CD45.2 or WT C57BL/6J mice (eight weeks old males) were irradiated in two doses of 550 rads each (for a total of 1,100 rads) four hours apart (RS-2000 X-Ray irradiator, Rad Source). Bone marrow cells from both femurs and tibias of donor mice (CFTR^Δ508^ male and CD45.1 wild-type male, eleven weeks old) were collected under sterile conditions. Bones were centrifuged for the collection of marrow, unfractionated bone marrow cells were washed, resuspended in PBS, mixed at a ratio of 1:1 or 1:2, confirmed by flow cytometry and injected retro-orbitally into the lethally irradiated mice (one donor to five recipients). Recipient mice were housed in a barrier facility under pathogen-free conditions before and after bone marrow transplantation. After bone marrow transplantation, mice were provided autoclaved acidified water with antibiotics (trimethoprim-sulfamethoxazole) and were fed autoclaved food. Mice were used for further experiments eight weeks after bone marrow reconstitution.

### Hematology

Blood counts of WT/Het or CFTR^Δ508^ littermates were taken via retro-orbital bleeding and analyzed by an automatic analyzer (Hemavet 950FS, DREW Scientific). Mice were anesthetized by the inhalation of isoflurane/oxygen gas mixture during the bleeding.

### Flow cytometry

Blood was obtained by cardiac puncture with an EDTA-coated syringe. Red blood cells were lysed in RBC Lysis Buffer according to the manufacturer’s protocol. Lungs were lavaged with PBS containing 2mM EDTA, dissociated by gentleMACS™ Dissociator (Miltenyi), and filtered through a 70 μm strainer. Ten centimeters of small intestines from the end of stomach, which contains duodenum and a part of jejunum, were collected. For preparation of lamina propria (LP) cells, fat and connective tissue were removed from intestines. Intestines were opened longitudinally, cut into 1-cm pieces and washed 3×10 minutes in HBSS containing 5% FBS, 2 mM EDTA, 100 mM N-acetyl cysteine, and 10 mM HEPES to remove mucus and epithelial cells. Tissue was digested with 1 mg/mL collagenase VIII (Sigma) and 20 μg/mL DNase I (Sigma) for 20 minutes at 37°C. Digested material was dissociated by gentleMACS™ Dissociator (Miltenyi) and filtered through a 70 μm strainer. Bone marrow cells were collected from both femurs and tibias. In some experiments, mice were challenged intranasally with mouse CCL2 100ng/mouse 12 hours prior to the harvest. Blood was collected via retro-orbital bleeding. BAL was collected by intratracheal lavage with PBS before the collection of lung.

All samples were collected in PBS with 2 mM EDTA to prevent cation-dependent cell-cell adhesion, and were stored on ice during transportation, staining and analysis. Cells were resuspended in 100 μl flow staining buffer (1% BSA and 0.1% sodium azide in PBS). Cells were stained with Ghost Dye™ Violet 510 for analysis of viability. Fcγ receptors were blocked for 15 min and surface antigens on cells were stained for 30 min at 4°C with directly conjugated fluorescent antibodies (flow cytometry antibody panels, Table S2). Forward-and side-scatter parameters were used for exclusion of doublets from analysis. Cell fluorescence was assessed with a LSRII (BD Biosciences) and was analyzed with FlowJo (BD, version 10.4). Gating scheme was shown in fig. S1–3. Lung and LP monocytes or macrophages were normalized by blood monocytes to eliminate the reconstructive rate difference of WT and CFTR^Δ508^ BM after BMT.

### Histology

Intestines were rinsed, opened on the anti-mesenteric side, cut into three strips, and placed in parallel on biopsy pad in a cassette. Samples were then fixed in zinc formalin, embedded in paraffin, and cut into 3- to 5-μm sections. The tissues were stained with hematoxylin and eosin (H&E) and periodic acid-Schiff (PAS) for histological assessment by a single investigator who was blinded to the experimental design. Slides were digitized on AxioScan Z1 slide scanner using 40× 0.95NA objective (Zeiss). The height of intestinal villi and crypt and the percentage of intestinal crypts that contain mucus was quantified in ZEN Blue software (Zeiss).

### Microfluidic perfusion assay

The assembly of the microfluidic devices used in this study and the coating of coverslips with recombinant mouse P-selectin-Fc and ICAM-1-Fc has been described previously *(37, 54–56).* Briefly, cleaned coverslips were coated with P-selectin-Fc (2 μg ml^−1^) and ICAM-1-Fc (10 μg ml^−1^) for 2 hours and then blocked for 1 hour with casein (1%) at RT. After coating, coverslips were sealed to polydimethylsiloxane (PDMS) chips by magnetic clamps to create flow chamber channels ~29 μm high and ~300 μm across. By modulating the pressure between the inlet well and the outlet reservoir, 6 dyn cm^−2^ wall shear stress was applied in all experiments.

For the in vitro adhesion assay of monocytes, isolated mouse BM cells (10^7^ cells ml^−1^) from WT or CFTR^ΔF508^ mice were incubated with PE-conjugated anti-CD115 antibody and AF647-conjugated anti-Ly6G antibody for 10 minutes at RT and perfused through the microfluidic device over a substrate of recombinant mouse P-selectin-Fc and recombinant mouse ICAM-1-Fc at a wall shear stress of 6 dyn cm^−2^. After cells were rolling on the substrate, 100 ng ml^−1^ mouse CCL2 was perfused. The processes of rolling and arrest were recorded by epifluorescence microscopy using an Olympus IX71 inverted microscope equipped with a 40 × 0. 95NA objective.

### Statistics

Statistical analysis was performed with Prism 6 (GraphPad). Data are presented as survival curve (Fig. 1A,3A,4A) and mean ± SD (Fig. 1B-F,I,J,2B,C,E,3C,D,F,4C-E,5B-G). Survival curves were compared using the Gehan-Breslow-Wilcoxon test. The means for the data sets were compared using the student t-tests with equal variances. P values less than 0.05 were considered significant.

## Supplementary Materials

Fig. S1. The gating scheme of blood flow cytometry assay.

Fig. S2. The gating scheme of lung cytometry assay.

Fig. S3. The gating scheme of LPL cytometry assay.

Table S1. Reconstitution rate of monocytes after BMT.

Table S2. Panels of flow cytometry assay.

## Acknowledgments

We thank Angela Denn from the Microscopy and Histology Core Facility at the La Jolla Institute for Allergy and Immunology for her help in obtaining scientific data presented in this paper. We thank Dr. Paul Quinton from the Department of Pediatrics in the School of Medicine at the University of California San Diego for his help in editing the manuscript. We thank Dr. Jesus Rivera-Nieves from the Inflammatory Bowel Disease Center in the Division of Gastroenterology at the University of California San Diego for his advice on histology.

## Author Contributions

Experiments were designed by Z.F. and K.L. Most experiments were performed by Z.F., J.M., R.H., E.E. and Z.M. Data analysis was performed by Z.F., Y.P.Z., and P.M.M. The manuscript was written by K.L., Z.F, and D.C. The project was supervised by K.L., and C.C.H. All authors discussed the results and commented on the manuscript.

## Data availability

The data that support the findings of this study are available from the corresponding author upon request.

## Supplementary Materials

Supplementary Figures

**Fig. S1.**
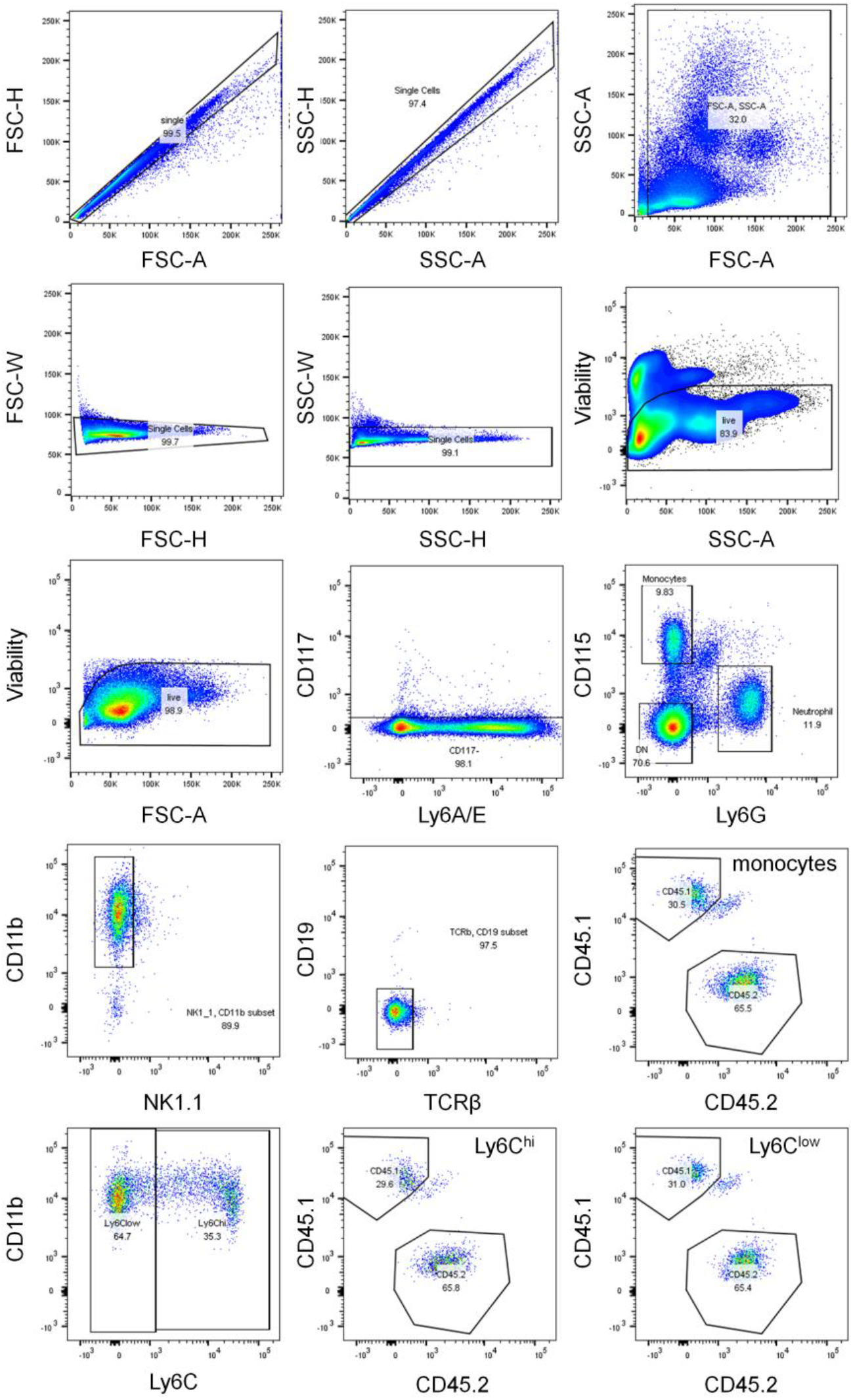
The gating scheme of blood flow cytometry assay. Singlets and live cells were gated first. Mature cells were gated as CD117^−^. Monocytes were gated as CD115^+^, Ly6G^−^, CD11b^+^, NK1.1^−^, CD19^−^, TCRβ^−^. Monocytes were identified as Ly6C^hi^ classical monocytes and Ly6C^low^ patrolling monocytes. In the mixed BMT mice, WT or CFTR^ΔF508^ monocytes were CD45.1+ or CD45.2+, respectively.

**Fig. S2.**
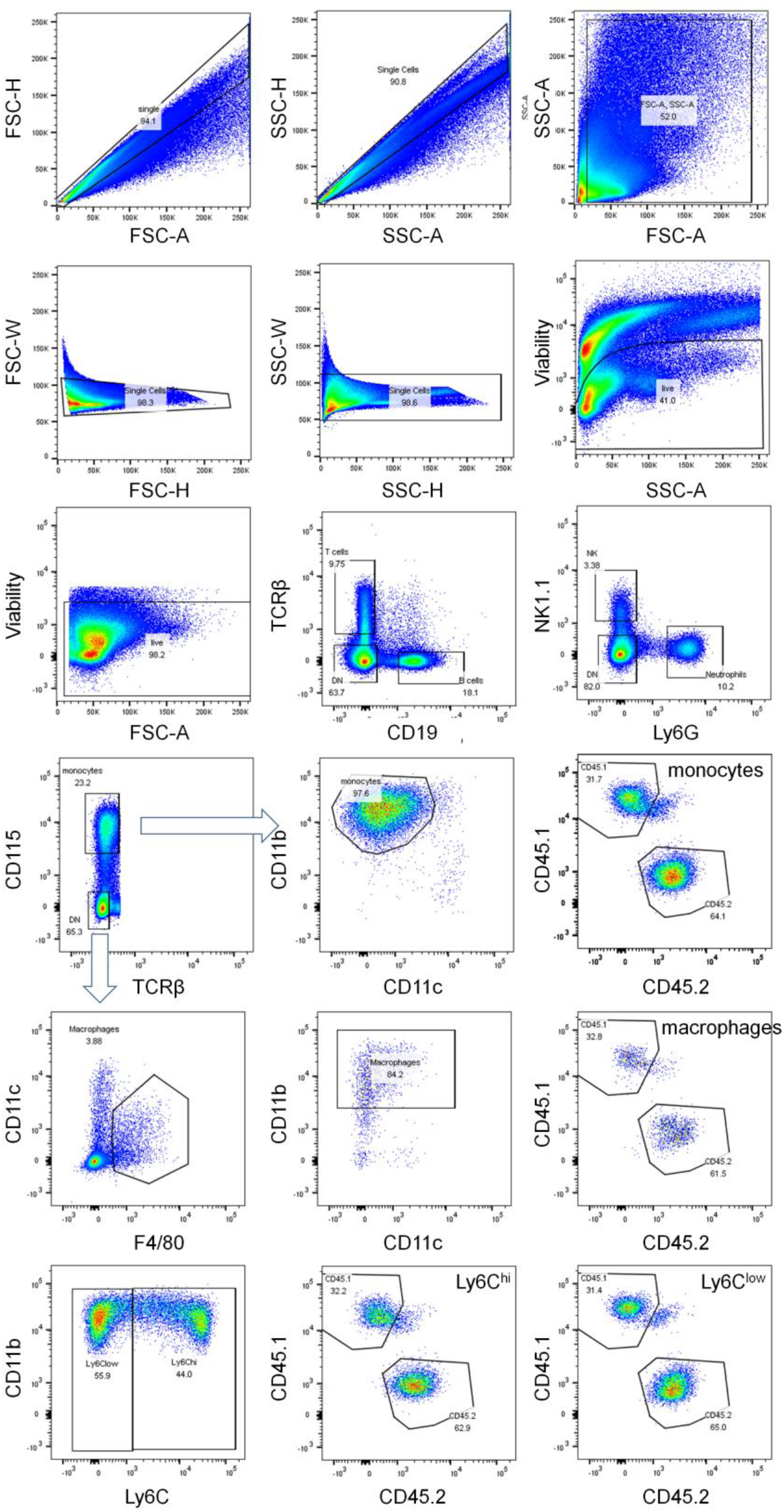
The gating scheme of lung cytometry assay. Singlets and live cells were gated first. Monocytes were gated as CD19^−^, TCRβ^−^, Ly6G^−^, NK1.1^−^, CD115^+^, CD11b^+^, CD11c^−^. Macrophages were gated as CD19^−^, TCRβ^−^, Ly6G^−^, NK1.1^−^, CD115^−^, F4/80+, CD11b^+^. Monocytes were identified as Ly6C^hi^ classical monocytes and Ly6C^low^ patrolling monocytes. In the mixed BMT mice, WT or CFTR^ΔF508^ monocytes were CD45.1+ or CD45.2+, respectively. The BAL gating is the same as lung gating.

**Fig. S3.**
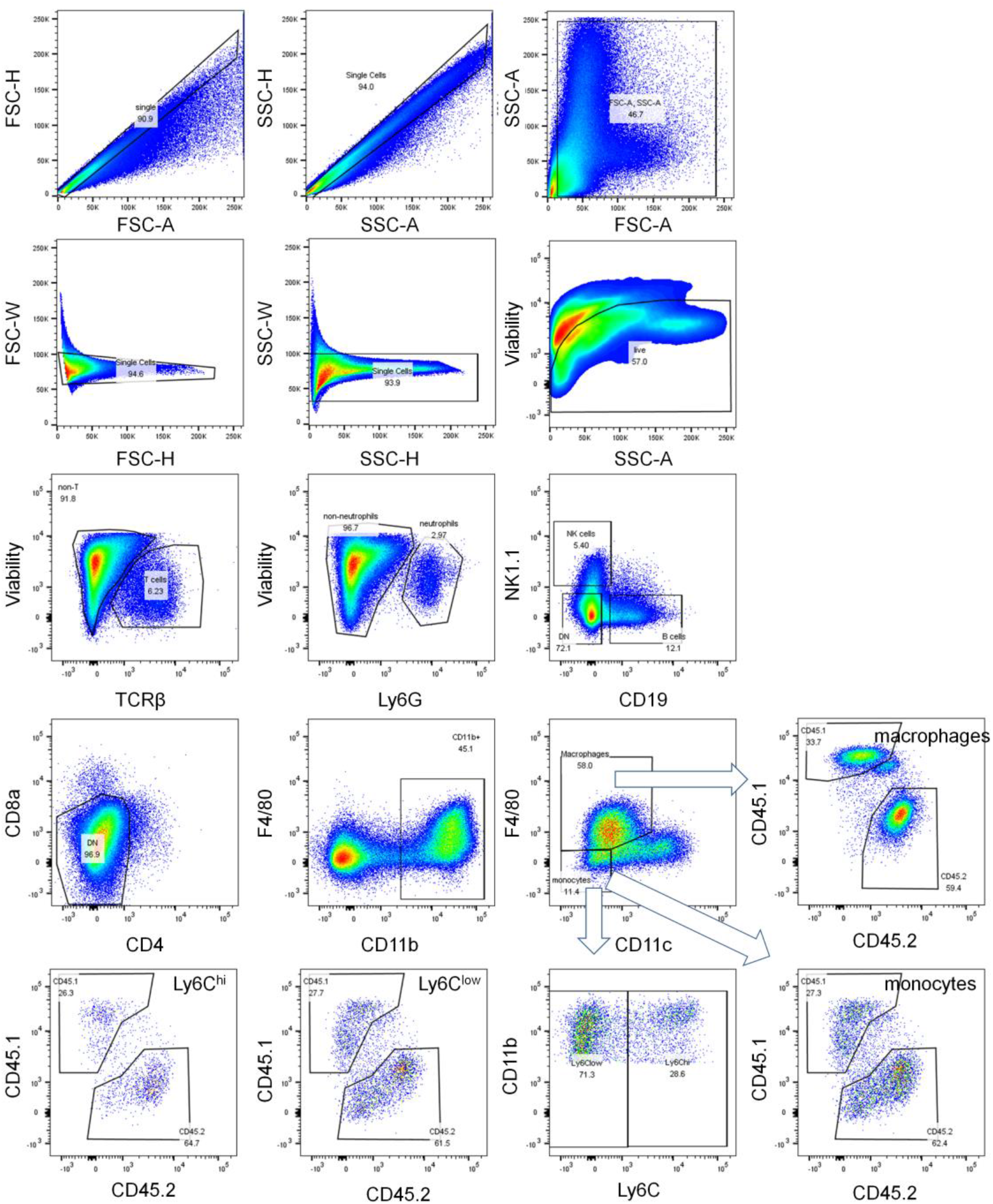
The gating scheme of LPL cytometry assay. Singlets and live cells were gated first. Monocytes were gated as TCRβ^−^, Ly6G^−^, CD19^−^, NK1.1^−^, CD4^−^, CD8a^−^, CD11b^+^, F4/80^−^, CD11c^−^. Macrophages were gated as TCRβ^−^, Ly6G^−^, CD19^−^, NK1.1^−^, CD4^−^, CD8a^−^, CD11b^+^, F4/80+. Monocytes were identified as Ly6C^hi^ classical monocytes and Ly6C^low^ patrolling monocytes. In the mixed BMT mice, WT or CFTR^ΔF508^ monocytes were CD45.1+ or CD45.2+, respectively.

**Supplementary Tables**

**Table S1.** Reconstitution rate of monocytes after BMT.

|  | WT to KO (700 rads) | WT to WT (1100 rads) | KO to WT (1100 rads) |
| --- | --- | --- | --- |
| BM | $67 \pm 2\%$ | $99.8 \pm 0.1\%$ | $99 \pm 1\%$ |
| Blood | $78 \pm 15\%$ | $99.9 \pm 0.0\%$ | $98 \pm 2\%$ |
| Lung | $80 \pm 13\%$ | $99.05 \pm 0.6\%$ | $97 \pm 3\%$ |
| Intestinal lamina propria | $60 \pm 6\%$ | $89 \pm 1\%$ | $86 \pm 4\%$ |

**Table S2.** Panels of flow cytometry assay.

| Antigen-clone | Color | Dilution |
| --- | --- | --- |
| Panel 1 (Blood, BM) |  |  |
| Viability | Ghost Dye™ Violet 510 | 1/1000 |
| CD45.1 -A20 | FITC | 1/200 |
| CD45.2 - 104 | PerCP-Cy5.5 | 1/200 |
| CD11b -M1/70 | BV785 | 1/200 |
| CD11c - N418 | BV605 | 1/200 |
| CD115 -AFS98 | PE-Cy7 | 1/100 |
| Ly6C -HK1.4 | BV570 | 1/200 |
| Ly6G - 1A8 | AF700 | 1/200 |
| TCRb - H57-597 | BV711 | 1/200 |
| CD4 - GK1.5 | PE-eFluor610 | 1/200 |
| CD8a - 53-6.7 | eFluor450 | 1/200 |
| CD19 - 6D5 | APC-Cy7 | 1/200 |
| CD117 (ckit) - 2B8 | APC | 1/100 |
| Ly6A/E(sca1) - D7 | PE | 1/100 |
| NK1.1 - PK136 | BV650 | 1/200 |
| Panel 2 (BAL, Lung, LPL) |  |  |
| Viability | BV510 | 1/1000 |
| CD45.1 -A20 | FITC | 1/200 |
| CD45.2 - 104 | PerCP-Cy5.5 | 1/200 |
| CD11b -M1/70 | BV785 | 1/200 |
| CD11c - N418 | BV605 | 1/200 |
| CD115 -AFS98 | PE-Cy7 | 1/100 |
| Ly6C -HK1.4 | BV570 | 1/200 |
| Ly6G - 1A8 | AF700 | 1/200 |
| TCRb - H57-597 | BV711 | 1/200 |
| CD4 - GK1.5 | PE-eFluor610 | 1/200 |
| CD8a - 53-6.7 | e450 | 1/200 |
| CD19 - 6D5 | APC-Cy7 | 1/200 |
| F4/80 - BM8 | PE | 1/100 |
| NK1.1 - PK136 | BV650 | 1/200 |

